# *N*-Aldehyde-Modified Phosphatidylethanolamines generated by lipid peroxidation are robust substrates of *N*-Acyl Phosphatidylethanolamine Phospholipase D

**DOI:** 10.1101/2024.10.30.621135

**Authors:** Reza Fadaei, Annie C. Bernstein, Andrew N. Jenkins, Allison G. Pickens, Jonah E. Zarrow, Abdul-Musawwir Alli-Oluwafuyi, Keri A. Tallman, Sean S. Davies

## Abstract

*N*-acyl phosphatidylethanolamine-hydrolyzing phospholipase D (NAPE-PLD) hydrolyzes phosphatidylethanolamines (PE) where the headgroup nitrogen has been enzymatically modified with acyl chains of four carbons or longer (*N*-acyl-PEs or NAPEs). The nitrogen headgroup of PE can also be non-enzymatically modified by reactive lipid aldehydes, thus forming *N*-aldehyde modified-PEs (NALPEs). Some NALPEs such as *N*-carboxyacyl-PEs are linked to PE via amide bonds similar to NAPEs, but others are linked by imine, pyrrole, or lactam moieties. Whether NAPE-PLD can hydrolyze NALPEs was unknown. We therefore characterized the major NALPE species formed during lipid peroxidation of arachidonic acid and linoleic acid and generated various NALPEs for characterization of their sensitivity to NAPE-PLD hydrolysis by reacting synthesized aldehydes with PE. We found that NAPE-PLD could act on NALPEs of various lengths and linkage types including those derived from PE modified by malondialdehyde (*N*-MDA-PE), butane dialdehyde (*N*-BDA-PE), 4-hydroxynonenal (*N*-HNE-PE), 4-oxo-nonenal (*N*-ONE-PE), 9-keto-12-oxo-dodecenoic acid (*N*-KODA-PE), and 15-E_2_-isolevuglandin (*N*-IsoLG-PE). To assess the relative preference of NAPE-PLD for various NALPEs versus its canonical NAPE substrates, we generated a substrate mixture containing roughly equimolar concentrations of the seven NALPEs as well as two NAPEs (*N*-palmitoyl-PE and *N*-linoleoyl-PE) and measured their rate of hydrolysis. Several NALPE species, including the *N*-HNE-PE pyrrole species, were hydrolyzed at a similar rate as *N*-linoleoyl-PE and many of the other NALPEs showed intermediate rates of hydrolysis. These results significantly expand the substrate repertoire of NAPE-PLD and suggest that it may play an important role in clearing products of lipid peroxidation in addition to its established role in the biosynthesis of *N*-acyl-ethanolamines.

## INTRODUCTION

Lipid peroxidation of polyunsaturated fatty acids (PUFAs) including linoleic acid and arachidonic acid generates a variety of lipid aldehydes that react with primary amines to form stable adducts [1]. These include malondialdehyde (MDA), butane dialdehyde (BDA, also known as succinaldehyde), 4-oxo-2-nonenal (ONE), 4-hydroxynonenal (HNE), 9-keto-12-oxo-dodecenoic acid (KODA), and isolevuglandins (IsoLG) [2-6]. MDA, HNE, and IsoLG have previously been shown to react with phosphatidylethanolamine (PE) to form *N*-aldehyde modified-PEs (NALPEs) [7]. Peroxidation of arachidonic acid in the presence of PE or of HDL also generates a series of *N*-carboxyacyl-PEs [8]. While the precise mechanism for the formation of *N*-carboxyacyl-PEs has not been fully elucidated, the most likely mechanism is by the reaction of PE with aldehyde species that retain the carboxylate moiety of the original PUFA during their formation by lipid peroxidation and where the initial reaction with PE forms an imine (Schiff base) bond that is subsequently oxidized to an amide bond by peroxides [8, 9].

Several previously characterized NALPEs have proinflammatory and cytotoxic effects [8]. Understanding how these NALPEs are degraded could lead to a better understanding of how inflammation in regulated. We hypothesized that *N*-acyl phosphatidylethanolamine-hydrolyzing phospholipase D (NAPE-PLD) may be an important enzyme for degrading NALPEs. NAPE-PLD is a zinc metallo-β-lactamase that catalyzes the hydrolysis of PEs that have been enzymatically modified on the headgroup nitrogen with acyl chains of four carbons or longer (NAPEs)[10, 11]. NAPE-PLD’s substrate preferences contrast with those of phospholipase D1 and D2, which are serine hydrolase-type enzymes that act on unmodified PEs and phosphatidylcholines, but not on NAPEs [12-14]. Homologs of NAPE-PLD are found in yeast, worms, reptiles, and mammals, likely indicating it has highly conserved functions in physiology [12, 15-17]. Our hypothesis that NAPE-PLD acts on substrates such as NALPEs was based on three previous observations. First, that the putative substrate binding site of NAPE-PLD is a large hydrophobic nook with several hydrophobic channels emanating from the active site that could potentially accommodate a wide variety of structures [18]. Second, that fluorogenic probes used to monitor NAPE-PLD activity in high throughput screening assays (i.e. PEDA1, PED6, and flame-NAPE) [19], all share a six-carbon *N*-acyl chain with an ω-dinitrophenyl moiety, a structure that shares some similarity to several previously reported *N*-carboxyacyl-PE species of NALPEs (Fig. 1). Third, that PE where the headgroup nitrogen is modified by the aldehyde 15-E_2_-isolevuglandin (*N*-IsoLG-PE) is a substrate for NAPE-PLD [20].

**Figure 1.**
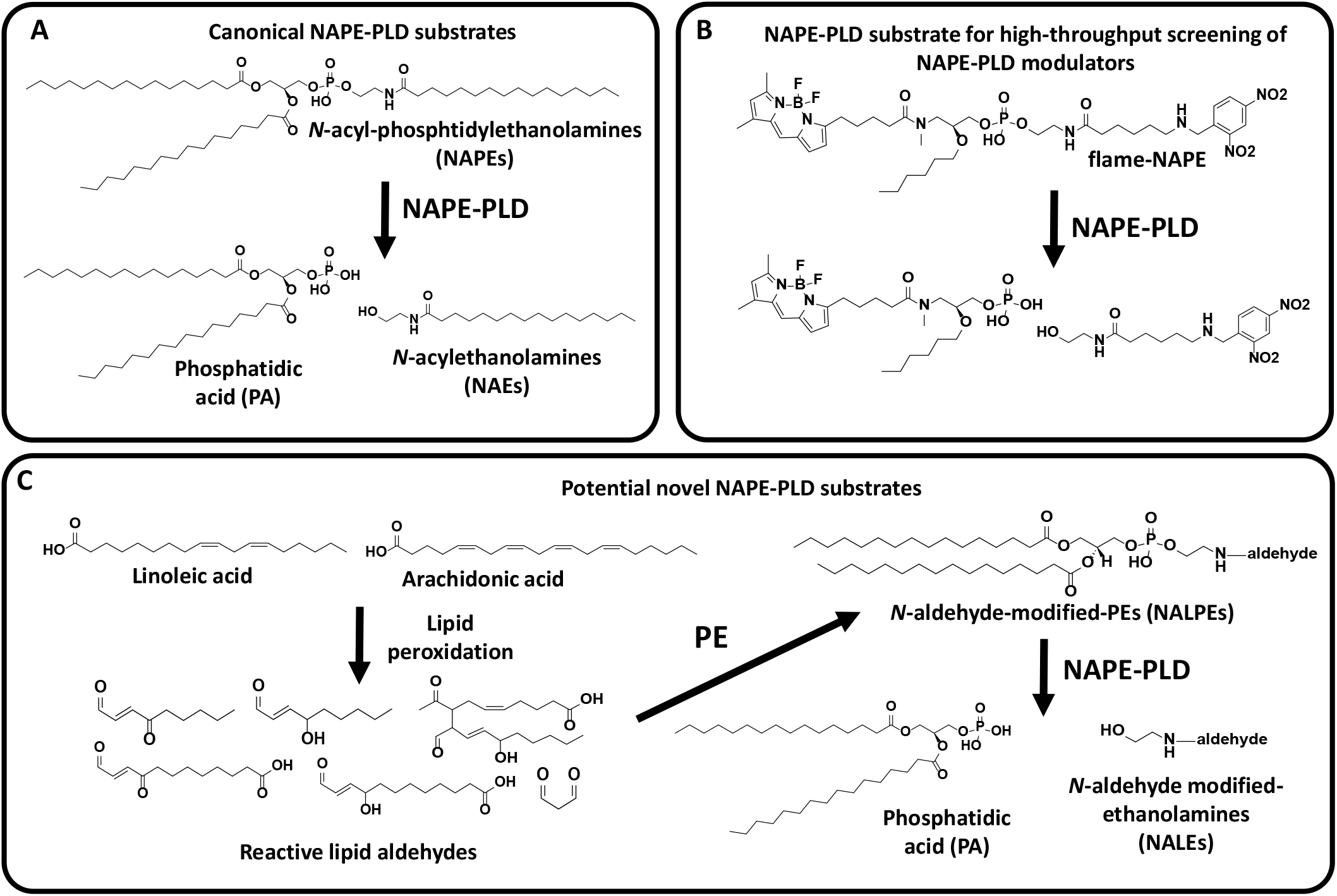
Canonical and hypothesized NAPE-PLD substrates. A) NAPE-PLD hydrolyzes its canonical substrates, *N*-acyl-phosphatidylethanolamines (NAPEs), to phosphatidic acid (PA) and *N*-acyl-ethanolamines. B) High-throughput screening for modulators of NAPE-PLD activity utilize fluorogenic probes such as flame-NAPE (pictured), PED6, and PED-A1, which have an *N*-acyl-nitrophenyl group as the quencher moiety to suppress BODIPY fluorescence. NAPE-PLD efficiently hydrolyzes these substrates to BODIPY-phosphatidic acid and *N*-acyl-nitrophenyl-ethanolamine suggesting that NAPE-PLD readily accommodates polar groups at the ω-terminus of the *N*-acyl chain. C) Lipid peroxidation of PUFAs including linoleic acid and arachidonic acid leads to formation of reactive aldehydes such as MDA, ONE, HNE, KODA, and HODA, and IsoLG, and these reactive aldehydes can react with the headgroup nitrogen of PE to form *N*-aldehyde modified-PEs (NALPEs). The present study tests whether these NALPEs are also efficient substrates of NAPE-PLD.

To test our hypothesis that NAPE-PLD efficiently hydrolyzes NALPEs in addition to its canonical substrates (e.g. NAPEs), we further characterized the major species of NALPEs generated by lipid peroxidation and then tested various synthetic NALPEs for their ability to be hydrolyzed by recombinant NAPE-PLD. We found that many different species of NALPEs are hydrolyzed by NAPE-PLD, some with equivalent efficiency as canonical NAPEs. These studies suggest that NAPE-PLD may be an important degradative enzyme of NALPEs in vivo, so that its reduced expression during cardiometabolic diseases might result in markedly increased NALPE levels and thereby contribute to inflammation.

## MATERIALS AND METHODS

### Materials and Reagents

1,2-Dipalmitoyl-sn-glycero-3-phosphoethanolamine (dpPE/PE), 1,2-dihexaoyl-sn-glycero-3-phosphoethanolamine (dhPE), *N*-glutaryl-PE, *N*-dodecanoyl-PE (also known as *N*-carboxyundecanoyl-PE (*N*-CUDA-PE)) were purchased from Avanti Polar Lipids (Alabaster, AL). *N*-Palmitoyl-(dipalmitoyl)PE (*N*-palmitoyl-PE) and *N*-linoleoyl-(dipalmitoyl)PE (*N*-linoleoyl-PE) were purchased from Echelon Biosciences (Salt Lake City, UT). Linoleic acid and arachidonic acid were purchased from Cayman Chemical, Michigan, USA. 1,1,3,3-Tetramethoxypropane (TMOP) was purchased from Sigma Aldrich and 2,5-dimethyltetrahydrofuran (DMTF) was purchased from Thermo scientific. Organic solvents, including methanol, chloroform, dichloromethane, and acetonitrile, were high-performance liquid chromatography grade from EMD Millipore.

### Oxidation of arachidonic acid/linoleic acid in the presence of PE

To investigate potential modifications of PE by reactive aldehydes resulting from lipid peroxidation, arachidonic acid and linoleic acid (10 mM each) were oxidized using 0.3 mM V70 as a free radical initiator in the presence of 0.5 mM PE in a reaction buffer of Triethylamine Acetate and Isopropanol (1:1 ratio), incubating at 37°C overnight with ample oxygen. The lipid was extracted using the 2:1 chloroform/methanol (v/v), dried under nitrogen gas,and dissolved in mobile phase A for liquid chromatography-mass spectrometry (LC/MS) injection. Two control samples were prepared exactly as the first except control one lacked PE and control two lacked arachidonic acid and linoleic acid. Another set of three samples was also prepared where dhPE was substituted for dpPE.

### Synthesis of *N*-aldehyde modified-phosphatidylethanolamines

IsoLG, KODA, ONE, and HNE were synthesized as previously described [21, 22]. BDA and MDA were freshly prepared by acidic conversion of DMTF and TMOP, respectively, as previously described [23]. Individual NALPEs were generated by reaction of aldehyde with dpPE overnight in reaction buffer (triethylamine acetate/isopropanol 1:1, v/v). The molar ratio of each aldehyde to PE was based on pilot studies to determine the amount of aldehyde needed to deplete >50% of PE. Final reactions used the following molar equivalents (ME) of aldehyde to PE: 3:1 BDA, 16:1 MDA; 10:1 HNE, 3:1 ONE, 5:1 KODA, 2:1 IsoLG. Reactions were extracted with 2:1 chloroform/methanol (v/v) and the yield of each product estimated based on LC/MS, with total yield based on reduction in PE and individual yield of individual species based on peak heights.

### Liquid Chromatography/Mass Spectrometry analysis of NALPEs

LC/MS analyses were performed using ThermoFinnigan Quantum Electrospray ionization triple-quadrupole mass spectrometer equipped with a surveyor autosampler operating in positive mode. For analysis of products from the reaction of aldehydes with dpPE, product mixtures were injected and separated on gradient high-performance liquid chromatography (HPLC) using a 50 × 2.1 mm C18 Kinetix column (2.6 um) directly coupled to the mass spectrometer with a constant flow rate of 0.1 ml/min. The mobile phase A consisted of a mixture containing isopropanol, methanol, and water (in a ratio of 5:1:4, v/v/v), enriched with 0.2% formic acid, 0.66 mM ammonium formate, and 0.003% phosphoric acid. Mobile phase B was composed of isopropanol with 0.2% formic acid. Starting condition were 5% mobile phase B and held for 0.5-min, followed by a gradient ramp to 95% mobile phase B over 2 minutes, which was then held for another 2 min before a rapid transition back to 5% mobile phase B over 0.5 min and the column allowed to reequilibrate for 2 min prior to the next injection. Electrospray ionization was performed at 3300 V, and the ion-transfer tube operated at 25 V and 270°C. The tube lens voltage was set to -180 V. Precursor scanning for reactions with dpPE was performed from *m/z* 350 to 1500 Da using product ion m/z 551.5 and for reactions with dhPE was performed from *m/z* 150 to 1200 Da using product ion m/z 271.2..

### Expression and purification of NAPE-PLD

Recombinant mouse NAPE-PLD was expressed in *Escherichia coli* as previously described with slight modifications [20]. The purification process involved lysis of induced bacterial cultures in lysis buffer (50 mM NaH_2_PO_4_, 300 mM NaCl, 10 mM imidazole, pH 8.0), incubation with TALON Superflow agarose beads contain cobalt ion, elution using elution buffer (50 mM NaH_2_PO_4_, 300 mM NaCl, 250 mM imidazole, pH 8.0), and washing with wash buffer (50 mM NaH2PO4, 300 mM NaCl, 20 mM imidazole, pH 8.0). Purified fractions were concentrated, dialyzed against buffer (Hanks’ Balanced Salt Solution), and stored at -80°C.

### Hydrolysis of *N*-aldehyde modified PEs by NAPE-PLD

To test the ability of NAPE-PLD to hydrolyze individual NALPEs, a reaction solution was prepared with the following reagents: 30 µM NALPE, 0.4% *N*-octyl glucoside, 50 mM Tris-HCl, and 20 ng/mL NAPE-PLD. The reaction was performed for each NALPE separately by incubation at 37°C for 2 hours and was quenched by extraction using a 2:1 chloroform/methanol mixture containing dioleoyl-phosphatidic acid (doPA) and d_4_-*N*-palmitoyl-(dioleoyl)PE (d_4_-NPPE) as internal standards. The organic layer was isolated and evaporated under nitrogen, followed by reconstitution in mobile phase A for analysis by LC/MS analysis using the instrument in multiple reaction monitoring (MRM) mode (Table 1). To compare the relative rate of hydrolysis of NAPE-PLD substrates directly, approximately equal concentrations of *N*-MDA-PE, *N*-BDA-PE, *N*-HNE-PE, *N*-ONE-PE, *N*-CUDA-PE, *N*-KODA-PE, *N*-IsoLG-PE, *N*-palmitoyl-PE, and *N*-linoleoyl-PE were combined together to create a substrate mixture. This substrate mixture was incubated with active enzyme and aliquots withdrawn after one, two and three hours of incubation. As a control, separate aliquots of the mixture were also incubated with heat-inactivated enzyme and processed in a similar manner. Samples incubated with inactive enzyme were used to establish the starting levels of each lipid without hydrolysis (0 h with active enzyme).

**Table 1.** Ions used for multiple reaction monitoring to quantify NAPE-PLD hydrolysis.

|  |  |  |
| --- | --- | --- |
| **NALPEs** | **m/z [M+H]<sup>+</sup> → m/z 551.5** | **m/z [M+NH<sub>4</sub>]<sup>+</sup> → m/z 551.5** |
| *N*-BDA-PE | 742.5 (pyrrole) | 759.5 (pyrrole) |
| *N*-MDA-PE | 746.5 (Schiff base) | 763.5 (Schiff base) |
| *N*-HNE-PE | 812.6 (pyrrole)<br>848.6 (Micheal adduct) | 829.6 (pyrrole)<br>865.6 (Micheal adduct) |
| *N*-ONE-PE | 828.6 (Schiff base)<br>846.6 (keto amide) | 845.6 (Schiff base)<br>863.6 (keto amide) |
| *N*-glutaryl-PE | 806.5 | 823.5 |
| *N*-CUDA-PE | 904.6 | 921.6 |
| *N*-KODA-PE | 900.6 (Schiff base)<br>918.6 (keto amide) | 917.6 (Schiff base)<br>935.6 (keto amide) |
| *N*-IsoLG-PE | 1006.7 (anhydrolactam)<br>1024.7 (lactam) | 1023.7 (anhydrolactam)<br>1041.7 (lactam) |
| **NAPEs** | **m/z [M+H]<sup>+</sup> → m/z 551.5** | **m/z [M+NH<sub>4</sub>]<sup>+</sup> → m/z 551.5** |
| *N*-palmitoyl-PE | 930.7 | 947.7 |
| *N*-linoleoyl-PE | 954.7 | 971.7 |
| **Internal standards** | **m/z [M+H]<sup>+</sup> → m/z 603.5** | **m/z [M+NH<sub>4</sub>]<sup>+</sup> → m/z 603.5** |
| dioleoyl-PA | 701.5 | 718.5 |
| d<sub>4</sub>-NPPE | 986.6 | 1003.6 |

## RESULTS

### Identification of major NALPE species formed during lipid peroxidation in vitro

To identify the major NALPEs that could form during lipid peroxidation, we oxidized arachidonic acid and linoleic acid in the presence of the endogenous PE, dipalmitoyl PE (dpPE). To confirm these results, we performed the same reaction with the model PE, dihexanoyl PE (dhPE), which is also somewhat more water-soluble. The resulting products for both reactions were analyzed by LC/MS in positive ion mode using precursor scanning. For reactions with dpPE, the product ion was set to *m/z 551*, while for dhPE the product ion set to *m/z 271*. The spectrum of peaks generated by precursor scanning of the reaction with dpPE is shown in Figure 2, while that for the reaction with dhPE is shown in Supplementary Figure 1. For simplicity, resulting products are referenced by their mass shift (in atomic mass units, amu) from the original PE used in the reaction. NALPE species formed in reactions with both dpPE and dhPE are listed in Supplemental Table 1.

**Figure 2.**
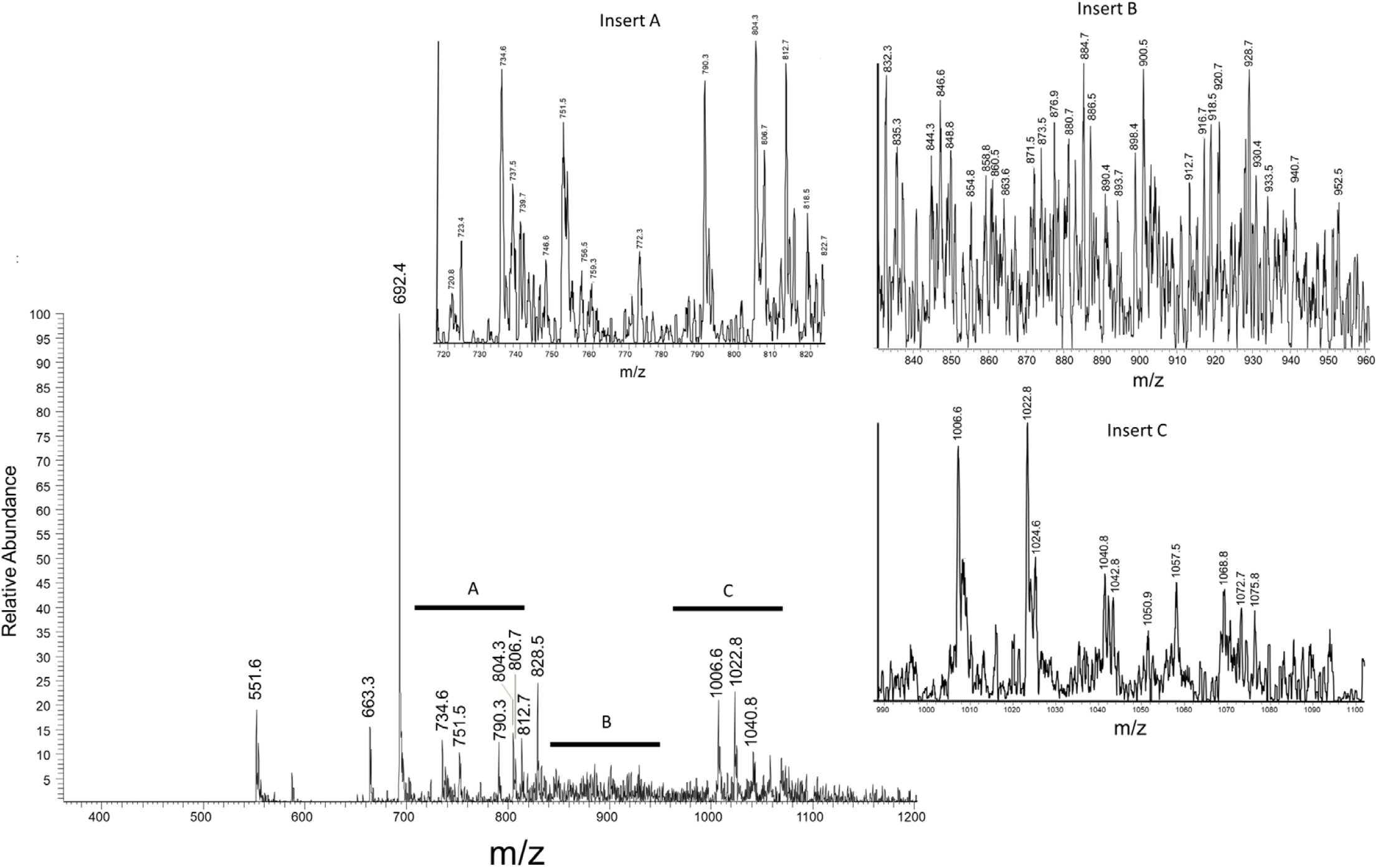
Identification of major NALPE species formed during lipid peroxidation. Arachidonic acid and linoleic acid were oxidized in the presence of dipalmitoyl-PE and the resulting products analyzed by LC/MS in positive ion mode using precursor scanning for product ion *m/z 551.5*. The mass spectrum of the broad peak seen in the total ion current chromatogram is shown, with inserts A, B and C providing magnified view of specific regions of the spectrum.

The largest peaks found in the reactions are consistent with aldehyde adduct species previously reported to form on PE or on other cellular primary amines such as lysine. The +42 amu NALPE and +59 amu NALPEs (*m/z 734.5* and *m/z 751.6* in Figure 2) are consistent with the [M+H]^+^ and [M+NH4]^+^ ions, respectively, of *N*-acetyl-PE. The +98 amu NALPE (*m/z 790.5* in Figure 2) and the +114 amu NALPE (*m/z 806.7*) are consistent with *N*-hexanoyl-PE and *N*-glutaryl-PE. *N*-hexanoyl and *N*-glutaryl adducts of lysine, 2-hydroxybenzylamine, and PE have previously been reported [8, 24, 25]. The +120 amu and +156 amu NALPEs (*m/z 812.8* and *m/z 848.6* in Figure 2) are consistent with the previously reported pyrrole and Michael addition adduct species of *N*-HNE-PE, respectively [26-28]. The +136 amu NALPE, along with the far more modest +154 amu NALPE species are consistent with the Schiff base and ketoamide species of *N*-ONE-PE. We note that it was previously reported that the Schiff base adduct of *N*-ONE-PE does not mature into a pyrrole adduct [26]. The +136 amu and +154 NALPEs could also represent the pyrrole and Schiff Base adduct species, respectively, arising from PE modification by 5-hydroxy-8-oxo-octenoic acid (HOOA). The esterified form of HOOA has been reported as a major form of oxidized phospholipid arising from the oxidation of 1-palmitoyl-2-arachidonyl-phosphatidylcholine (PAPC) [29]. The +314, +330, +332, and +348 amu NALPEs (*m/z 1006.8, 1022.7, 1024.6* and *1040.7*, respectively, in Figure 2) are consistent with the anhydrolactam, anhydrohydroxylactam, lactam, and hydroxylactam species, respectively, of *N*-IsoLG-PE. These adduct species have been previously reported for the reaction of isolevuglandin or oxidized arachidonic acid with lysine, 2-hydroxybenzylamine, and PE [21, 30].

Several of the more modest peaks are also consistent with those previously reported or predicted to arise from known lipid aldehydes generated during peroxidation. The +28 amu and +45 amu NALPEs are consistent with the [M+H]^+^ and [M+NH_4_]^+^ ions of *N*-formyl-PE, which could arise from formaldehyde reaction with PE. The +54 amu NALPE (peak 746.5) is consistent with the Schiff Base product formed by the reaction of malondialdehyde (MDA) with PE (*N*-MDA-PE) and was identified to form in oxidized HDL [8]. The +208 amu and the +226 amu NALPEs are consistent with the mass expected from the reaction of 9-keto-12-oxo-dodecenoic acid (KODA) with PE to form Schiff base and ketoamide adducts, respectively. The +192 amu NALPE peak (*m/z 884.7*) is consistent with the mass expected from the reaction of 9-hydroxy-12-oxo-dodecenoic acid (HODA) with PE to form a pyrrole adduct. The esterified forms of KODA and HODA has been reported as a major form of oxidized phospholipid arising from the oxidation of 1-palmitoyl-2-linoleoyl-phosphatidylcholine (PLPC) [29]. We could not readily identify predicted structures for other minor peaks detected as clusters of peaks between *m/z 850.6* and *m/z 1006.6*, although they could represent various *N*-carboxyacyl-PEs and other lower yield NALPEs.

### Synthesis of NALPEs for hydrolysis studies

To confirm the identity of the putative NALPE species (e.g. *N*-KODA-PE) and to test whether individual NALPEs are robust substrates for NAPE-PLD, we reacted dpPE with synthesized versions of major endogenous aldehydes or their analogs, characterized the major products and determined whether these products were substrates for hydrolysis by NAPE-PLD. We did not synthesize or test *N*-formyl-PE, *N*-acetyl-PE, and *N*-hexanoyl-PE, as these have previously been tested as substrates for NAPE-PLD, with only *N*-hexanoyl-PE found to be a substrate [19].

We considered the putative ketoamide adduct of *N*-ONE-PE as the most likely NALPE to be a substrate of NAPE-PLD, as it only differs from the canonical substrates of NAPE-PLD, NAPEs, by the addition of a ketone group at carbon 4. Reaction of ONE with dpPE yielded two major species, one consistent with the predicted Schiff Base adduct (*m/z 828.6*) and the other consistent with the predicted ketoamide adduct (*m/z 846.6*) (Figure 3A and supplemental figure 2A). Incubation of *N*-ONE-PE with active NAPE-PLD for two hours resulted in >90% reduction in signal for the *N*-ONE-PE ketoamide species and about a 65% reduction in signal for the *N*-ONE-PE Schiff base species, compared to the signal obtained when *N*-ONE-PE was incubated with heat-inactivated enzyme (Figure 3B and supplemental figure 2B). Incubation with active enzyme also significantly increased phosphatidic acid levels (Figure 3C and supplemental figure 2C).

**Figure 3.**
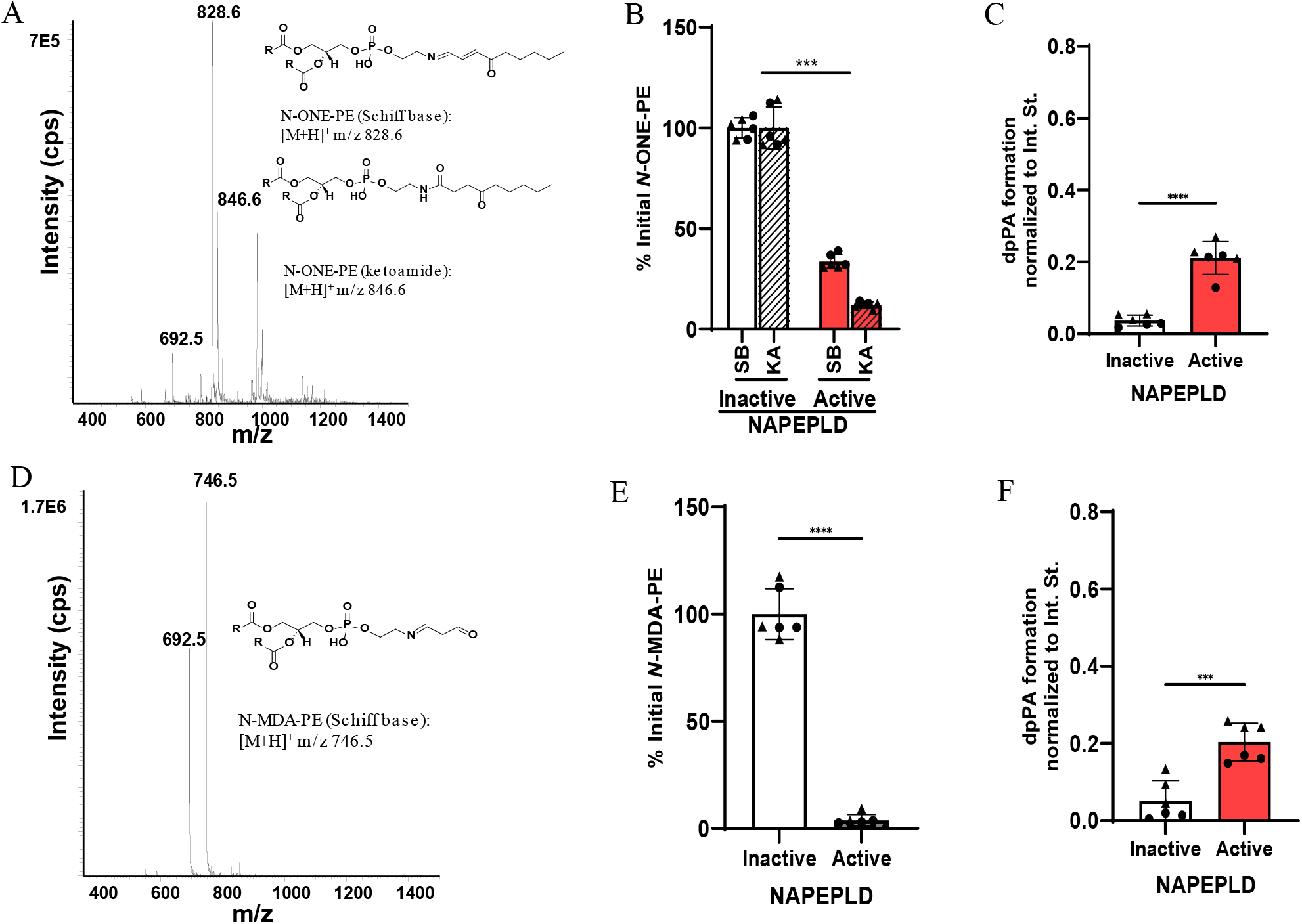
N-ONE-PE and *N*-MDA-PE are hydrolyzed by NAPE-PLD. A) Dipalmitoyl-PE ([M+H]+ m/z 692.5) was reacted with 4-oxo-nonenal (ONE) and the resulting products analyzed by LC/MS in positive ion mode using precursor scanning with m/z 551.5 product ion. The putative structure of the two major *N*-ONE-PE products are shown. Active recombinant NAPE-PLD or heat-inactivated enzyme was incubated with *N*-ONE-PE for 2 h and the changes in *N*-ONE-PE species (B) and phosphatidic acid (C) levels determined by LC/MS. The initial level of each *N*-ONE-PE species was set as the amount detected in the sample with inactive enzyme. Phosphatidic acid is shown as ratio compared to internal standard dioleoyl phosphatidic acid. D) Dipalmitoyl-PE was reacted with malondialdehyde (MDA) and the resulting *N*-MDA-PE products analyzed in a similar manner as *N*-ONE-PE. The putative structure of the major *N*-MDA-PE product is shown. *N*-MDA-PE was incubated with active or inactive NAPE-PLD for 2 h and the resulting change in levels of *N-*MDA-PE (E) and phosphatidic acid (F) monitored by LC/MS in a similar manner as for *N*-ONE-PE. For each, mean ± S.E.M. is shown with student’s t-test comparison of active vs inactive enzyme, ***p<0.001, and ****p<0.0001.

Given the prominent formation of a Schiff Base adduct with *N*-ONE-PE, we chose to further explore the effect of this alternative linkage compared to the amide bond on the ability of NAPE-PLD to hydrolyze substrate. Reaction of MDA with PE generated a product of *m/z 746.5*, consistent with the Schiff base species of *N*-MDA-PE, (Figure 3D and Supplemental Figure 2D). Incubation of *N*-MDA-PE with active NAPE-PLD for two hours resulted in essentially complete loss of the *N*-MDA-PE signal (Figure 3E and Supplemental Figure 2E), along with significant production of phosphatidic acid (Figure 3F and Supplemental Figure 2F). These results are consistent with NAPE-PLD being able to effectively utilize NALPEs with Schiff Base linkages as substrates.

HNE only differs from ONE by the addition of two hydrogens, but HNE is known to form distinct adducts from that of ONE. Reaction of HNE with PE yielded species consistent with the pyrrole (*m/z 812.6*) and Michael addition adducts (*m/z 848.6*) (Figure 4A and supplemental Figure 3A).Unlike most NALPEs which elute prior to PE on the reversed phase HPLC gradient used in our studies, the *N*-HNE-PE pyrrole species eluted after PE, consistent with being much more hydrophobic than other NALPEs. Incubation of *N*-HNE-PE with active NAPE-PLD for two hours resulted in essentially complete loss of the *N*-HNE-PE pyrrole signal, about 50% loss of the *N*-HNE-PE Michael adduct signal (Figure 4B and Supplemental Figure 3B), and significant formation of phosphatidic acid (Figure 4C and Supplemental Figure 3C).

**Figure 4.**
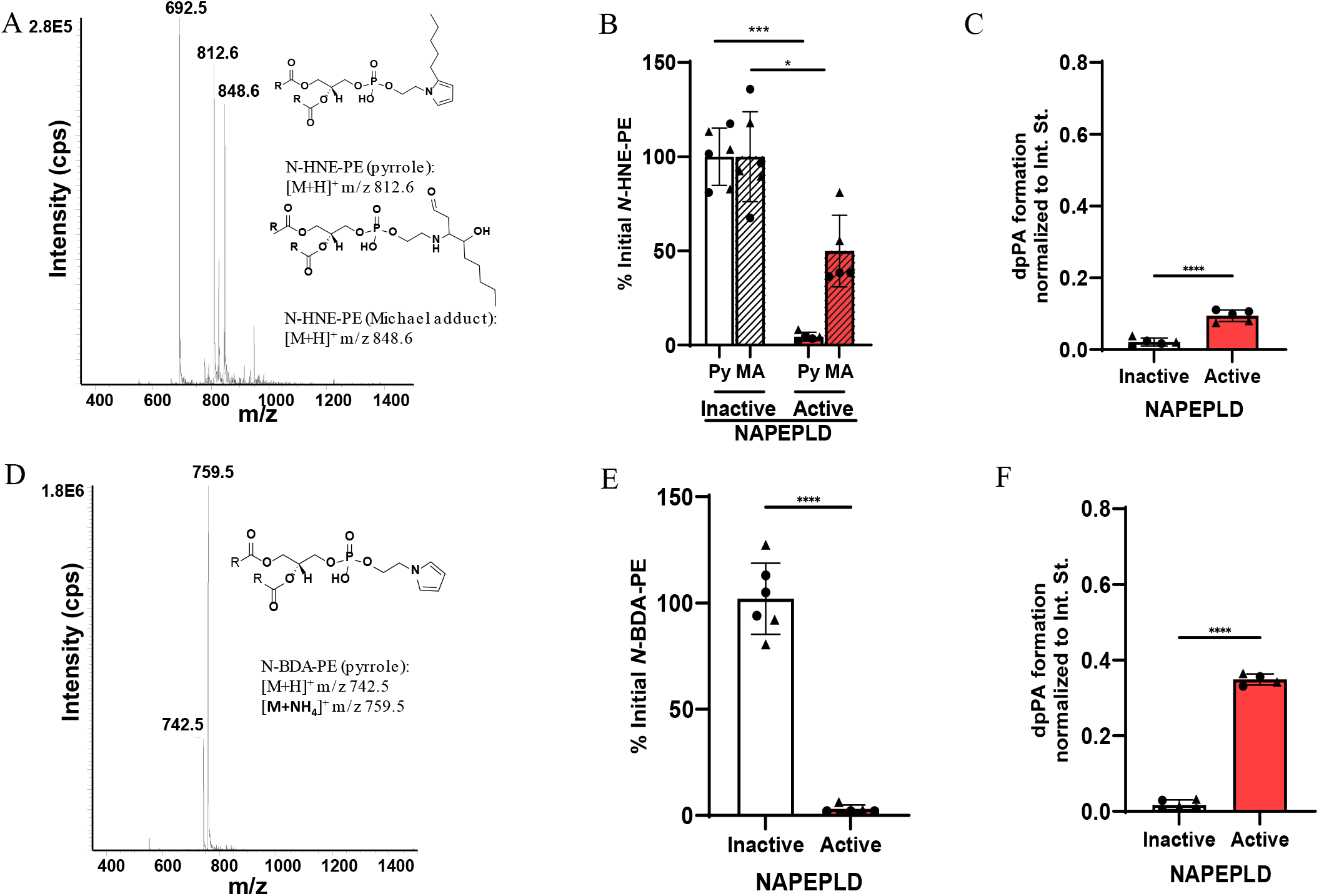
*N*-HNE-PE and *N*-BDA-PE are hydrolyzed by NAPEPLD. A) Dipalmitoyl-PE was reacted with 4-hydroxy-nonenal (HNE) and the resulting products analyzed by LC/MS in positive ion mode using precursor scanning with *m/z* 551.5 product ion. The putative structures of the two major *N*-HNE-PE products are shown. Active or heat-inactivated NAPE-PLD were incubated with *N*-HNE-PE for 2 h and changes in levels of *N*-HNE-PE (B) and phosphatidic acid (C) determined by LC/MS. D) Dipalmitoyl-PE was reacted with butane dialdehyde (BDA) and the resulting reactions products analyzed by LC/MS. The putative structure of the major *N*-BDA-PE product is shown. E, F) *N*-BDA-PE was incubated with active or inactive NAPE-PLD and the resulting changes in the levels of *N*-BDA-PE (E) and phosphatidic acid (F) measured by LC/MS. For each, mean ± S.E.M. is shown with student’s t-test comparison of active vs inactive enzyme, *p<0.05***p<0.001, and ****p<0.0001.

To further investigate the ability of NAPE-PLD to hydrolyze substrates with *N*-pyrrole linkages, we reacted butane dialdehyde (also known as succinaldehyde) with PE and characterized the resulting products. This reaction generated products of *m/z 742.5* and *m/z 759.5*, consistent with the [M+H]^+^ and [M+NH_4_]^+^ ions for the *N*-pyrrole species (Figure 4D and supplemental figure 4D). We note that similar *N*-pyrrole adducts of lysine and DNA have been previously reported [31], but we did not detect signals consistent with the *N*-BDA-PE pyrrole in our oxidation mixture. Incubation of *N*-BDA-PE with active NAPE-PLD resulted in essentially complete hydrolysis of this pyrrole species (Figure 4E and Supplemental Figure 3E) and significant formation of phosphatidic acid (Figure 4F and Supplemental Figure 3F).

In each of the above reactions, the hydrolyzed products had either an alkyl tail or were entirely hydrophobic. However, a number of NALPEs are *N*-carboxylacyl-PEs, i.e. they retain the terminal carboxylate moiety of the original fatty acid from which the lipid aldehyde is produced. We had previously found that NAPE-PLD could hydrolyze *N*-IsoLG-PE, which has such a terminal carboxylate. To confirm our previous results, we reacted 15-E_2_-isolevuglandin (IsoLG) with PE and characterized the resulting products. This reaction generated multiple adduct species, with the most prominent being those with *m/z 1006* and *m/z 1024*, consistent with the formation of the anhydrolactam and lactam species of *N*-IsoLG-PE, among others (Figure 5A and Supplemental Figure 4A). When we incubated this *N*-IsoLG-PE mixture with NAPE-PLD for two hours, we found that about 75% of both the anhydrolactam (AL) and lactam (Ltm) species were hydrolyzed (Figure 5B and Supplemental Figure 4B), with significant production of phosphatidic acid (Figure 5C and Supplemental Figure 4C).

**Figure 5.**
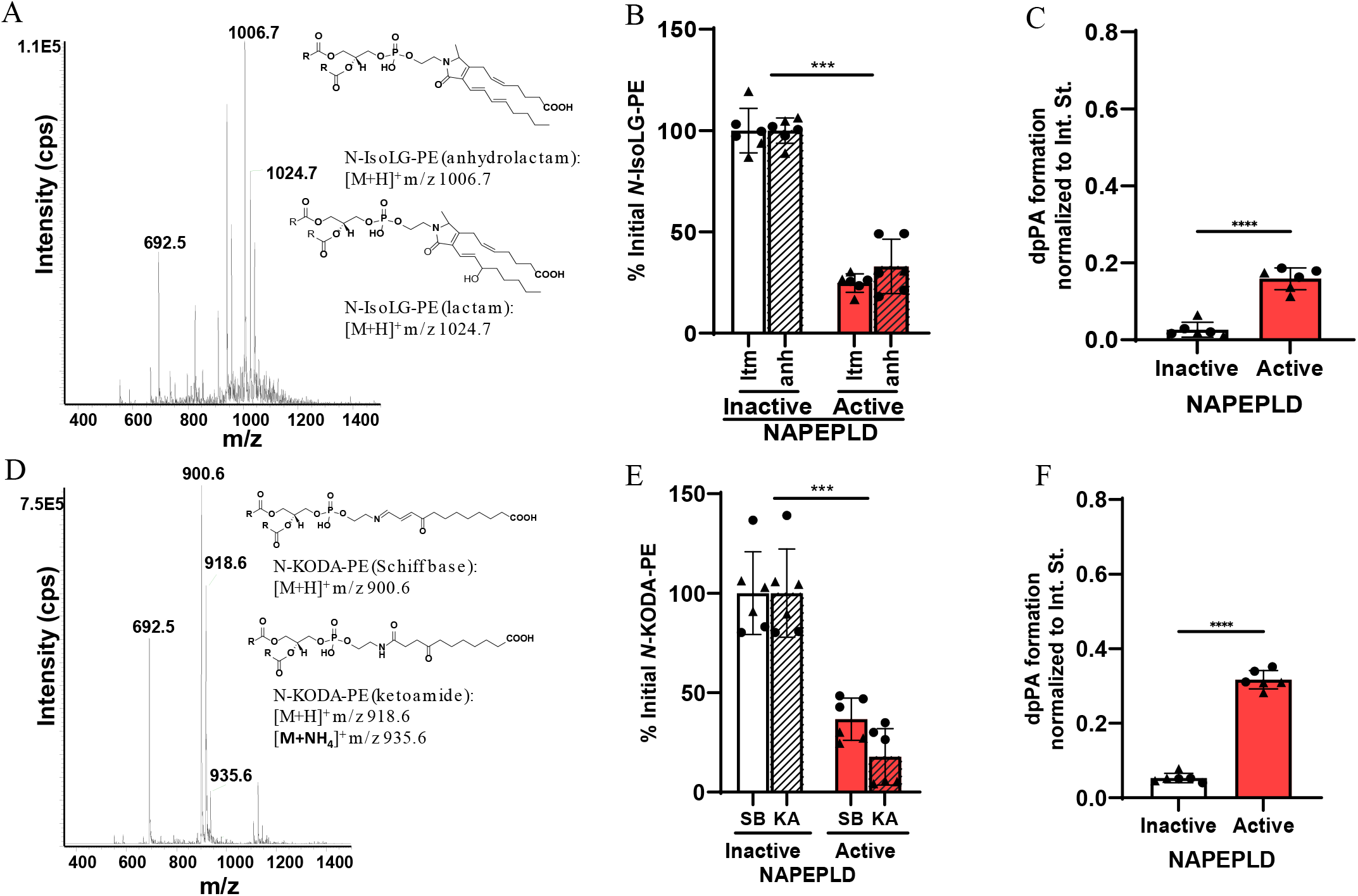
*N-*IsoLG-PE and *N*-KODA-PE are hydrolyzed by NAPEPLD. A) Dipalmitoyl-PE was reacted with IsoLG and the resulting products analyzed by LC/MS in positive ion mode using precursor scanning with *m/z* 551.5 product ion. The putative structures of the major *N*-IsoLG-PE products are shown. Active recombinant NAPE-PLD or heat inactivated enzyme was incubated with *N*-IsoLG-PE for 2 h, and the resulting changes in the levels of *N*-IsoLG-PE (B) and phosphatidic acid (C) measured by LC/MS. D) Dipalmitoyl-PE was reacted with 9-keto-12-oxo-dodecenoic acid (KODA) and the resulting products analyzed by LC/MS in a similar manner as for *N*-IsoLG-PE. Active or heat-inactivated NAPE-PLD was incubated with *N*-KODA-PE for 2 h and changes in levels of *N*-KODA-PE (E) and phosphatidic acid (F) determined by LC/MS. For each, mean ± S.E.M. is shown with student’s t-test comparison of active vs inactive enzyme, *p<0.05***p<0.001, and ****p<0.0001.

To further explore the effect of the terminal carboxylate moiety, we investigated whether *N*-KODA-PE is a substrate for NAPE-PLD. KODA is an analog of ONE that similarly results from the peroxidation of linoleic acid, except that it derives from fragmentation of the α-end of the molecule so that it retains the terminal carboxylate moiety. Reaction of KODA with PE yielded products whose masses were consistent with a Schiff Base adduct (*m/z 900.6*) and a ketoamide adduct (*m/z 918.6*) (Figure 5D and Supplemental Figure 4D). Incubation of *N*-KODA-PE with NAPE-PLD resulted in about 60% hydrolysis of the Schiff Base adduct and 80% hydrolysis of the ketoamide adduct (Figure 5E and Supplemental Figure 4E), with robust production of phosphatidic acid (Figure 5F and Supplemental Figure 4F).

To further investigate the effect of the terminal carboxylate, we purchased *N*-CUDA-PE and *N*-glutaryl-PE and confirmed these products by LC/MS (Figure 6 A,D and Supplemental Figure 5A,D). *N*-CUDA-PE differs from the *N*-KODA-PE ketoamide species only by the absence of the ketone group at carbon 4. Incubation of *N*-CUDA-PE with NAPE-PLD for 2 h resulted in its complete hydrolysis (Figure 6B and Supplemental Figure 5B) and robust increases in phosphatidic acid formation (Figure 6C and Supplemental Figure 5C). In contrast, incubation of *N*-glutaryl-PE with NAPE-PLD resulted in no hydrolysis (Figure 6E and Supplemental Figure 5E) or significant increases in phosphatidic acid formation (Figure 6F and Supplemental Figure 5F).

**Figure 6.**
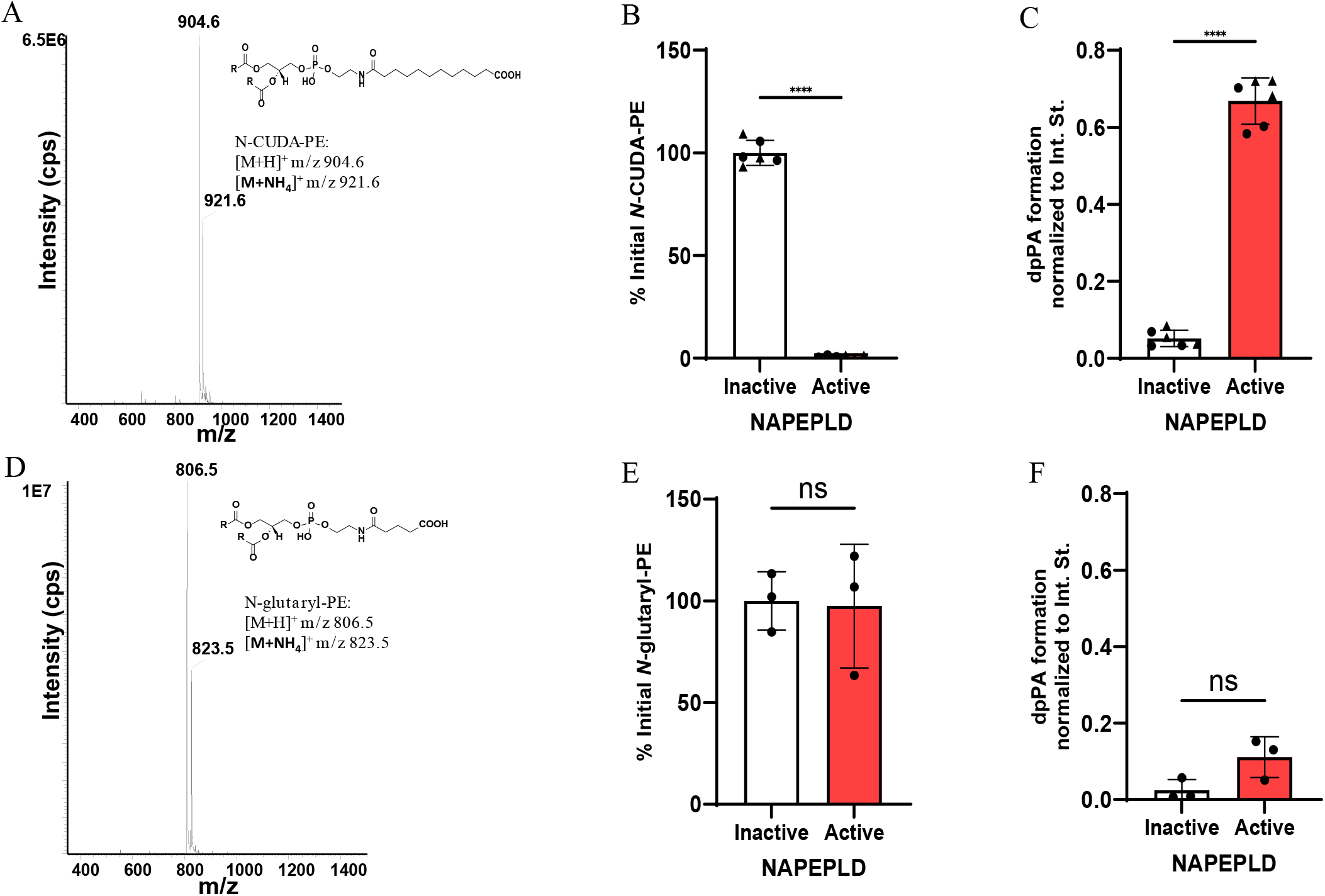
*N*-CUDA-PE and *N*-glutaryl-PE are hydrolyzed by NAPEPLD. A) *N*-CUDA-PE was purchased from a commercial source and analyzed by LC/MS in positive ion mode using precursor scanning with the product ion set to m/z 551. The putative structure of *N*-CUDA-PE is shown. Active or heat-inactivated NAPE-PLD was incubated with *N*-CUDA-PE for 2 h and changes in levels of *N*-CUDA-PE (B) and phosphatidic acid (C) determined by LC/MS. D) *N*-glutaryl-PE were purchased from a commercial source and analyzed by LC/MS in a similar manner as for *N*-CUDA-PE. Active or heat-inactivated NAPE-PLD was incubated with *N*-glutaryl-PE for 2 h and changes in levels of *N*-glutaryl-PE (E) and phosphatidic acid (F) determined by LC/MS. For each, mean ± S.E.M. is shown with student’s t-test comparison of active vs inactive enzyme, ns, non-significant, ***p<0.001, and ****p<0.0001.

### Comparing hydrolysis rate of NAPE-PLD substrates

During oxidative stress, NAPE-PLD is likely to encounter multiple possible substrates at the same time, so that the relative hydrolysis rate for each substrate will likely determine the extent to which each is degraded. To compare the relative rate that various NAPE-PLD substrates are hydrolyzed under competitive conditions, we created a substrate mixture by combining two NAPEs, *N*-palmitoyl-PE and *N*-linoleoyl-PE, with the various synthetic NALPEs, all in approximately equal amounts. We then incubated this substrate mixture with NAPE-PLD and monitored the progress of the reaction by observing the formation of phosphatidic acid (Figure 7A) and the loss of each starting substrates (Figure 7B-D). (For ease of visualization, results for the individual substrates within the mixture are separated into three graphs, with the same *N*-palmitoyl-PE curve shown in each for reference.)

**Figure 7:**
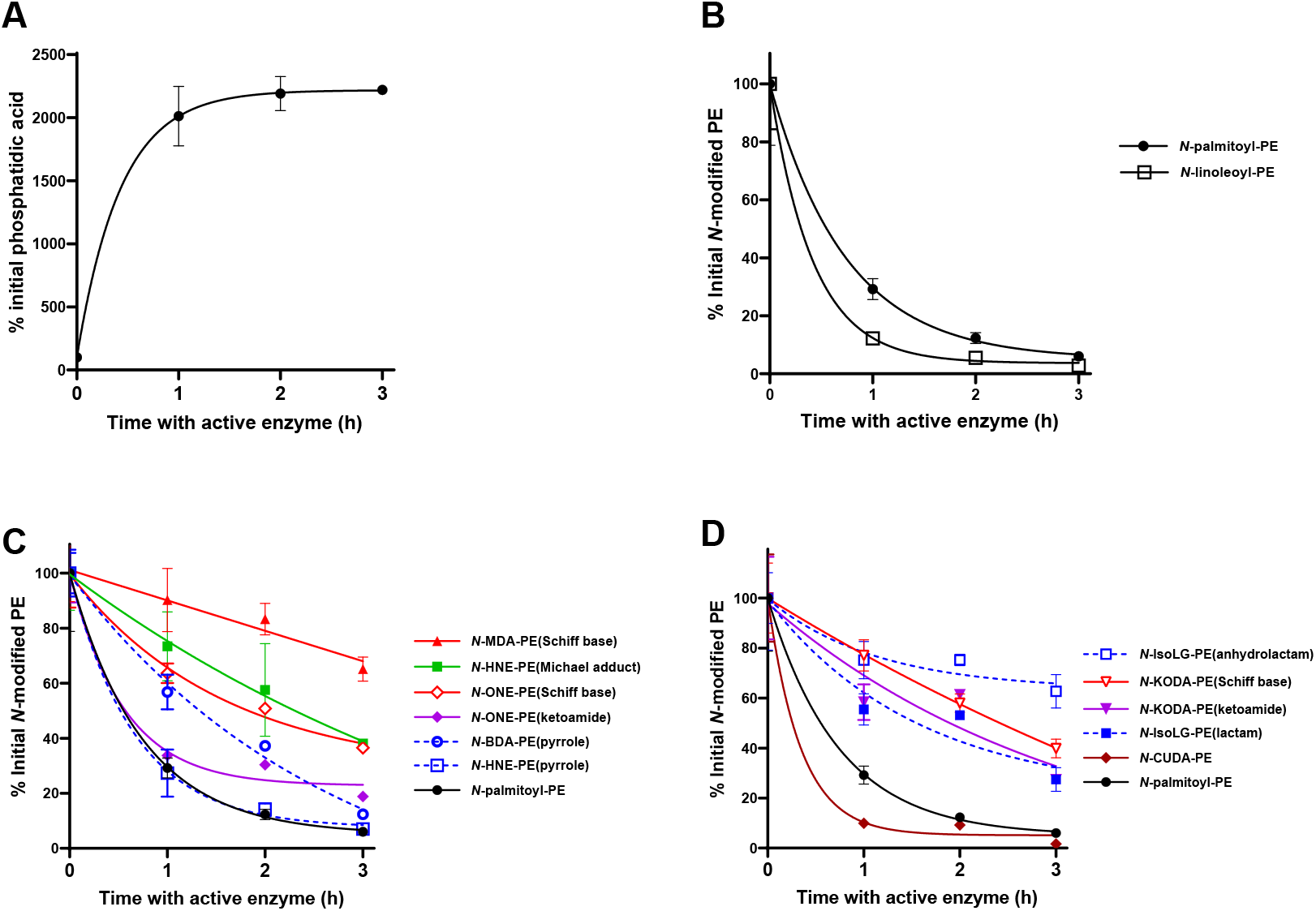
Individual species of NALPEs significantly vary in their relative rate of hydrolysis by NAPE-PLD. Active NAPE-PLD was incubated with a substrate mixture containing approximately equimolar amounts of *N*-palmitoyl-PE, *N*-linoleoyl-PE, *N*-MDA-PE, *N*-BDA-PE, *N*-HNE-PE, *N*-ONE-PE, *N-*KODA-PE, *N-*CUDA-PE, and *N*-IsoLG-PE and aliquots removed after 1, 2, and 3 h of incubation and analyzed by LC/MS. Levels of each starting lipid were normalized to values obtained when the substrate mixture was incubated with heat-inactivated NAPE-PLD (0 h incubation with active enzyme). A) Phosphatidic acid levels. B-D) For readability, the results are separated for NAPEs (B), short- and medium-chain NALPEs (C), and NALPEs species with carboxylate moiety (D) with the same *N*-palmitoyl-PE data shown on each graph for reference.

The two canonical NAPE-PLD substrates, *N*-palmitoyl-PE and *N*-linoleoyl-PE (Figure 7B), as well as *N*-HNE-PE (pyrrole) (Figure 7C), and *N*-CUDA-PE (Figure 7D) all exhibited similarly high rates of hydrolysis. *N*-BDA-PE(pyrrole) and *N*-ONE-PE(ketoamide) were hydrolyzed slightly slower (Figure 7C). *N*-KODA-PE(ketoamide) (Figure 7C) and *N*-IsoLG-PE(lactam) (Figure 7D) were hydrolyzed at a similar rate that was slightly slower than *N*-ONE-PE(ketoamide). *N*-ONE-PE(Schiff base) was hydrolyzed significantly slower than *N*-ONE-PE(ketoamide) (Figure 7C), and *N*-KODA-PE(Schiff Base) was hydrolyzed somewhat slower than *N*-KODA-PE(ketoamide) (Figure 7D). *N*-HNE-PE(Michael adduct) was hydrolyzed at a similar rate to *N-*ONE-PE(Schiff base) (Figure 7C). *N*-IsoLG-PE(anhydrolactam) and *N*-MDA-PE(Schiff base) were hydrolyzed at the slowest rates (Figure 7C and 7D).

## DISCUSSION

Previous research has documented the formation of NALPEs species both in vitro and in vivo [7-9, 27, 32-34], but little has been known about how these molecules are metabolized. In the studies reported here we demonstrate that many NALPEs are robust substrates for NAPE-PLD, thereby expanding the biological role of this enzyme to potentially include regulating the levels of these bioactive lipids in vivo. Our present study also identified KODA adducts of PE, which results from oxidation of linoleic acid, extending the total number of known NALPE species from previous studies that showed the formation of *N*-IsoLG-PE, *N*-MDA-PE, *N*-ONE-PE, *N*-HNE-PE, *N*-glutaryl-PE, *N*-formyl-PE, *N*-hexanoyl-PE, *N*-7:1carboxyacyl-PE, and *N*-9:2carboxyacyl-PE [7, 27, 32-39]. We had predicted that peroxidation of linoleic acid would lead to the formation of both Schiff Base and ketoamide PE adducts of KODA, based on previous studies that had characterized lysine and DNA adducts of KODA[40].

Our interest in identifying enzymes that metabolize various NALPEs arose from previous studies reporting significant biological activity for several NALPEs. For instance, *N*-HNE-PE and *N*-ONE-PE affect the function and activity of membrane proteins such as UCP1 by changing membrane fluidity and are suggested to affect other membrane proteins by a similar mechanism [41]. NALPEs such as *N*-MDA-PE, *N*-HNE-PE, *N*-ONE-PE and *N*-IsoLG-PE, but not *N*-glutaryl-PE, induce the pro-inflammatory responses of endothelial cells [8]. *N*-IsoLG-PEs also induces endoplasmic reticulum stress in endothelial cells [42]. *N*-IsoLG-PE also strongly activated NFκB and induced inflammatory cytokine expression in macrophages via RAGE [8, 43]. We had previously found that NAPE-PLD could hydrolyze *N*-IsoLG-PE to reduce its pro-inflammatory effects [20], and therefore hypothesized that NAPE-PLD might also hydrolyze other NALPEs.

Our studies provide structure activity relationships (SAR) for NAPE-PLD well beyond those previously established for NAPEs [44]. For instance, the presence of an ω-carboxylate group on the acyl chain of a NALPE (i.e. *N*-carboxyacyl-PEs) appears to have little detrimental effect on NAPE-PLD hydrolysis when the acyl chain is long but abolishes activity when the acyl chain is short (e.g. *N*-CUDA-PE is hydrolyzed as rapidly as either of the two tested NAPEs but *N*-glutaryl-PE are not hydrolyzed). One possible explanation for this would be if an extended acyl chain positions the polar ω-carboxylate group sufficiently far away from a hydrophobic channel in the active site that it only minimally disrupts hydrophobic interactions between the acyl carbons proximal to the amide bond and the hydrophobic channel of the enzyme. This notion is consistent with the finding that NALPEs with a ketone group three carbons from the amide bond (i.e. the ketoamide species) were somewhat less favorable substrates than NAPEs (e.g. the slower rate of hydrolysis for *N*-ONE-PE(ketoamide) than either NAPE or the slower rate of hydrolysis for the *N*-KODA-PE(ketoamide) compared to *N*-CUDA-PE, which only differ by the ketone on carbon 4). Another important SAR insight this study provides is the effect of alternative bonds linking the lipid chain to the PE nitrogen. NALPEs with *N*-pyrrole linkages were rapidly hydrolyzed, with the addition of an alkyl chain to the pyrrole further increasing the rate of hydrolysis (e.g. *N*-HNE-PE(pyrrole) vs *N*-BDA-PE(pyrrole)). In contrast, the Schiff base species of NALPEs such as *N*-MDA-PE, *N*-ONE-PE, and *N*-KODA-PE were poorly hydrolyzed when presented within a mixture that included other substrates. Finally, NALPEs that had two alkyl tails extending from the nitrogen linkage, such as the *N*-HNE-PE Michael adduct species and the various *N*-IsoLG-PE species, were hydrolyzed more slowly than those with a single tail in the competition experiments, suggesting that NAPE-PLD is somewhat less able to accommodate such substrates. Altogether, our findings that NAPE-PLD can robustly hydrolyze a wide variety of NALPEs suggest that NAPE-PLD might play an important role in regulating their levels *in vivo* and that some NALPEs may be more likely to accumulate due to slower degradation.

NAPE-PLD expression is reduced in inflammatory conditions including atherosclerosis, ulcerative colitis and neurodegenerative diseases [45-47]. The finding that that hepatocyte-specific deletion of *Napepld* renders mice more susceptible to inflammation [48] and adipocyte-specific deletion results in low-grade inflammation [49] suggests that the loss of NAPE-PLD has a causative role in inflammation.

Furthermore, bone marrow derived macrophages lacking NAPE-PLD have reduced capacity to carry out efferocytosis [50], a critical step in the resolution of inflammation. Our finding that NAPE-PLD robustly hydrolyzes pro-inflammatory NALPEs raises the possibility that some of the pro-inflammatory effects of reduced *Napepld* expression could arise from increases in the levels of NALPEs. Future studies are needed to assess whether loss of NAPE-PLD increases NALPE levels in various tissues and whether this contributes to the pro-inflammatory phenotype associated with loss of NAPE-PLD.

## Abbreviations

Anh: anhydrolactam
BDA: butane dialdehyde
CUDA: carboxyundecanoyl
DMTF: 2,5-dimethyltetrahydrofuran
doPA: dioleoyl-phosphatidic acid
dpPE: 1,2-dipalmitoyl-sn-glycero-3-phosphoethanolamine
d4-NPPE: d4-N-Palmitoyl-phosphatidylethanolamine
dhPE: 1,2-dihexaoyl-sn-glycero-3-phosphoethanolamine
glut: Glutaryl
HNE: 4-hydroxynonenal
HOOA: 5-hydroxy-8-oxo-octanoic acid
HPLC: high-performance liquid chromatography
IsoLG: 15-E2-isolevuglandin
KODA: 9-keto-12-oxo-dodecenoic acid
LCMS: Liquid chromatography-mass spectrometry
Ltm: lactam
MDA: malondialdehyde
NALPEs: N-aldehyde modified-phosphatidylethanolamines
NAPE-PLD: N-acyl phosphatidylethanolamine-hydrolyzing phospholipase D
NLPE: N-Linoleoyl-PE
NPPE: N-Palmitoyl-PE
ONE: 4-oxo-nonenal
PA: phosphatidic acid
PAPC: 1-palmitoyl-2-arachidonyl-phosphatidylcholine
PE: phosphatidylethanolamine
TMOP: 1,1,3,3-tetramethoxypropane.

## DATA AVAILABILITY STATEMENT

All data supporting the findings of this study are available within this article and its supplemental data.

## SUPPLEMENTAL DATA

This article contains supplemental data.

## AUTHORS CONTRIBUTIONS

**R.F**. assisted with conceptualization of the project, performed experiments, analysis, and visualization of results for publication, curated the data, co-wrote the original draft, and edited the final draft. **A.N.J**., **A.C.B**., **and A.G.P**. performed experiments and analysis and reviewed the final draft. **J.E.Z**. and **A.M.A.O**. generated, purified, and characterized recombinant NAPE-PLD and reviewed the final draft. **K.A.T**. synthesized and validated various lipid aldehydes and reviewed the final draft. **S.S.D**. conceptualized the project, acquired funding for and administered the project, supervised the overall project, assisted with analysis and visualization of the results, co-wrote the original draft, and edited the final draft.

