## Supplemental Information for "*N*-Aldehyde-Modified Phosphatidylethanolamines generated by lipid peroxidation are robust substrates of *N*-Acyl Phosphatidylethanolamine Phospholipase D"

[1]Department of Pharmacology, Vanderbilt University. Nashville, TN, USA, 37232; [2]College of Arts and Sciences, Vanderbilt University, Nashville, TN, USA; [3]Department of Cell Biology and Physiology, and [4]Department of Plant and Wildlife Sciences, Brigham Young University. Provo, UT, 84602; [5]Department of Chemistry, Vanderbilt University, Nashville, TN, USA; and [6]Vanderbilt Institute of Chemical Biology, Vanderbilt University, Nashville, TN, USA, 37235.

*To whom correspondence should be addressed.

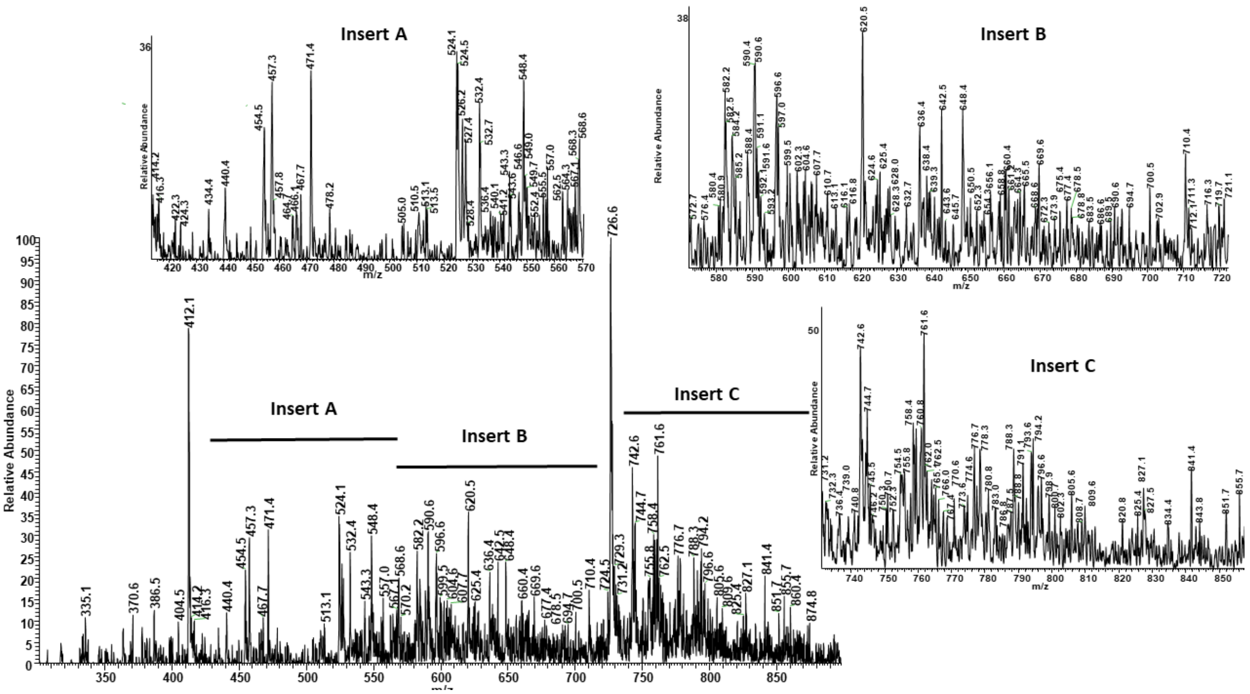


**Supplemental Figure1. Identification of major NALPE species formed during lipid peroxidation.** Arachidonic acid and linoleic acid were oxidized in the presence of dihexanoyl-PE. The resulting products were analyzed by LC/MS in positive ion mode using precursor scanning with *m/z* 271.2 as product ion. The mass spectrum of the major broad chromatographic peak is shown with magnified parts indicated by insert A, B and C. Full list of putative structure for significant peaks (peak height >3% peak height of PE) is provided in Supplemental Table 1.

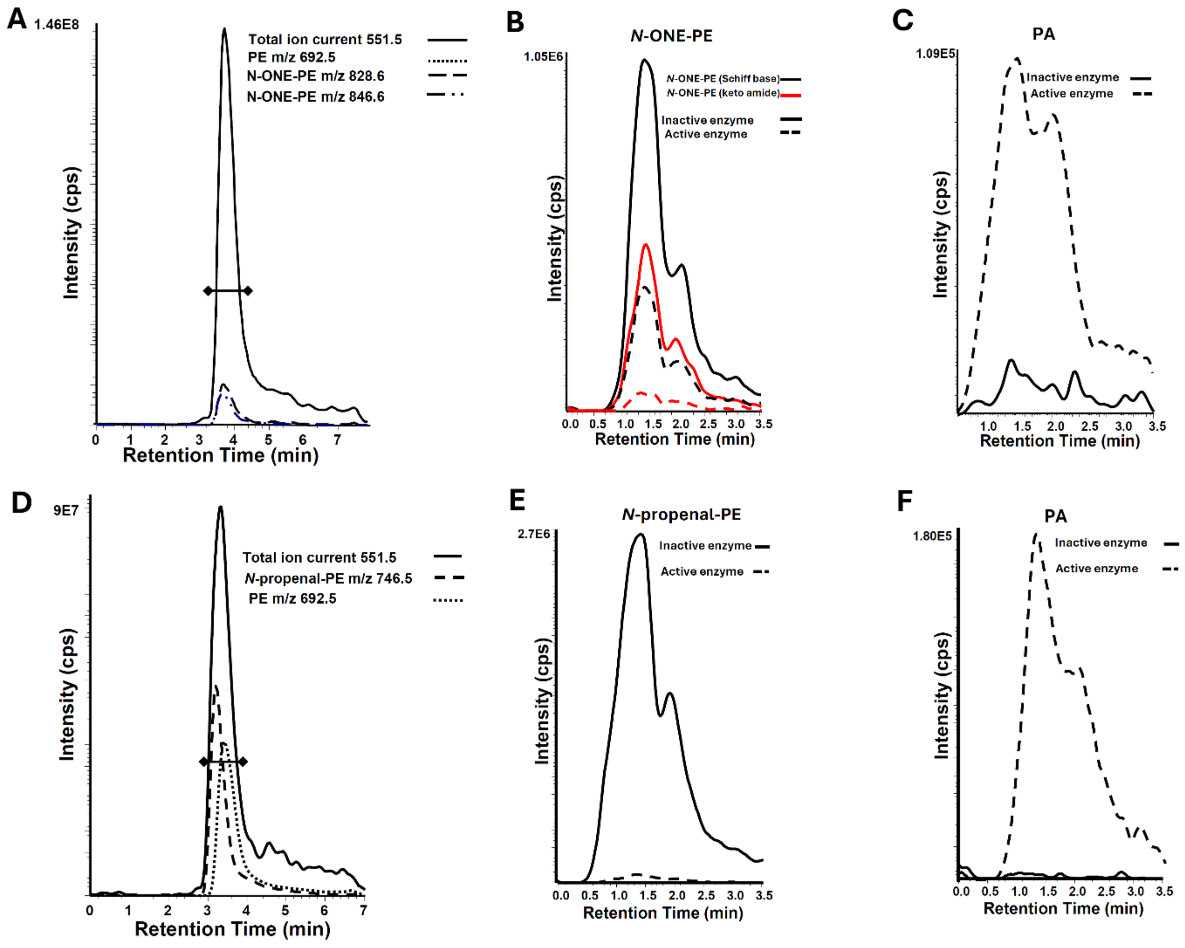


**Supplemental Figure 2. Chromatograms corresponding to LC/MS spectra and NAPEPLD hydrolysis of *N*-ONE-PE and *N*-MDA-PE in Figure 3.** A) Precursor scanning chromatogram (product ion *m/z 551.5*) of the reaction of ONE with PE, showing the formation of *N*-ONE-PE, including Schiff base and ketoamide species. Solid line represents the total ion current from precursor scan with retention time used for spectra shown in Figure 3A indicated by (♦—♦). Dotted lines represent reconstructed single ion current for major ions found in the spectrum showing *N*-ONE-PE species. B) LC/MS in MRM mode to monitor *N*-ONE-PE species and C) phosphatidic acid (PA) as product of the reaction. D) Precursor scanning chromatogram (product ion *m/z 551.5*) of the reaction of MDA with PE, resulting in the formation of N-MDA-PE Schiff base (aka *N*-propenal-PE). Solid line represents the total ion current of the precursor scan with retention time used for the spectra shown in Figure 3D indicated by (♦—♦). Dotted lines represent reconstructed single ion current for major ions found in the spectrum showing *N*-MDA-PE(Schiff base). E) LC/MS in MRM mode to monitor *N*-MDA-PE(Schiff base) and F) phosphatidic acid (PA) as product of the reaction.

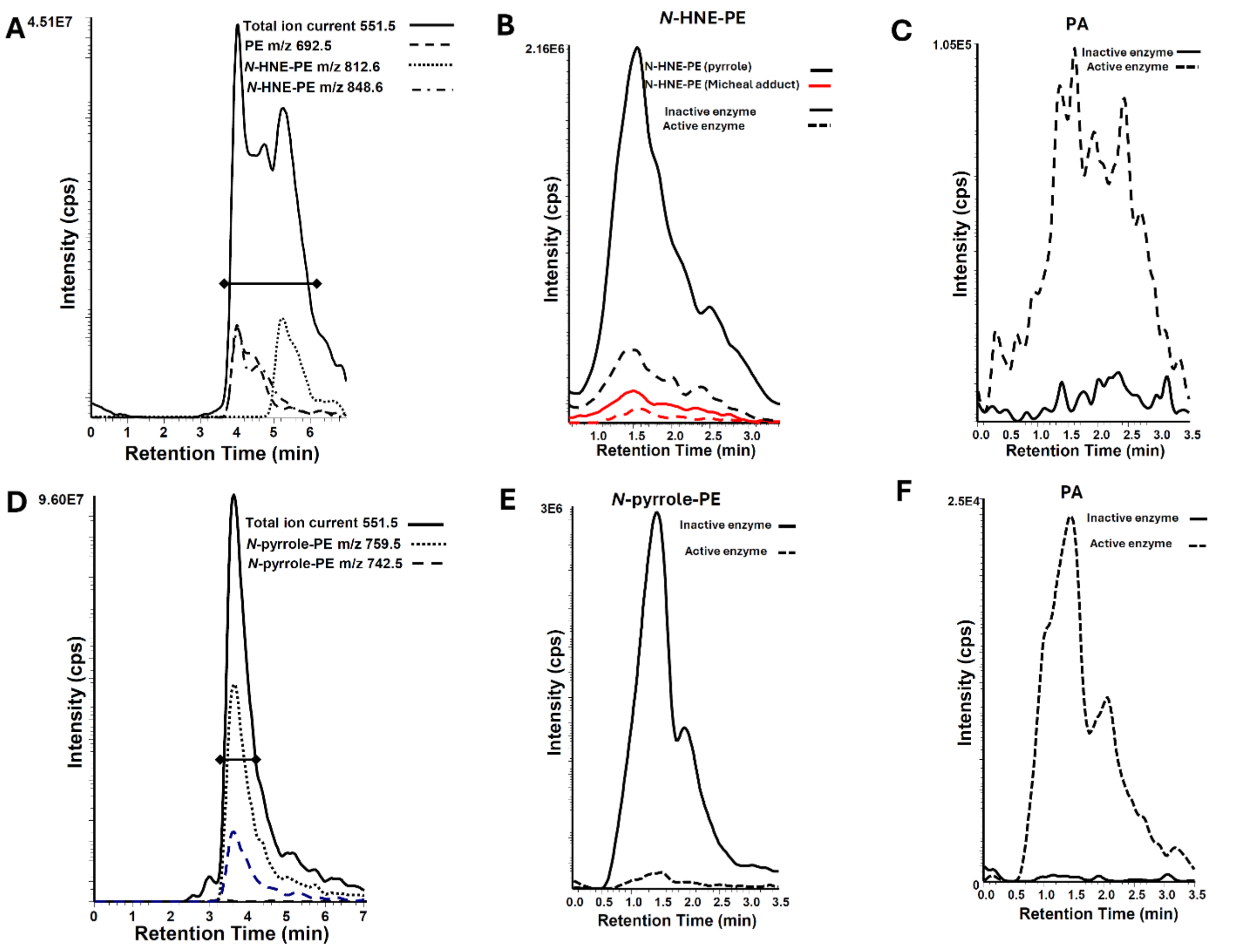


**Supplemental Figure 3. Chromatograms corresponding to LCMS spectra NAPEPLD hydrolysis of *N*-HNE-PE and *N*-BDA-PE in Figure 4.** A) Precursor scanning chromatogram (product ion *m/z 551.5*) of the reaction of HNE with PE, resulting in the formation of *N*-HNE-PE, including pyrrole and Michael adduct species. Solid line represents the total ion current from precursor scan with retention time used for spectra shown in Figure 4A indicated by (♦—♦). Dotted lines represent reconstructed single ion current for major ions found in the spectrum showing *N*-HNE-PE spices. B) LC/MS in MRM mode to monitor *N*-HNE-PE species and C) phosphatidic acid (PA) as product of the reaction. D) Precursor scanning chromatogram (product ion *m/z 551.5*) of the reaction of BDA with PE, resulting in the formation of *N*-BDA-PE(pyrrole) (aka *N*-pyrrole-PE). Solid line represents the total ion current from precursor scan with retention time used for spectra shown in Figure 4D indicated by (♦—♦). Dotted lines represent reconstructed single ion current for major ions found in the spectrum showing *N*-pyrrole-PE. E) LC/MS in MRM mode to monitor *N*-pyrrole-PE and F) phosphatidic acid (PA) as product of the reaction.

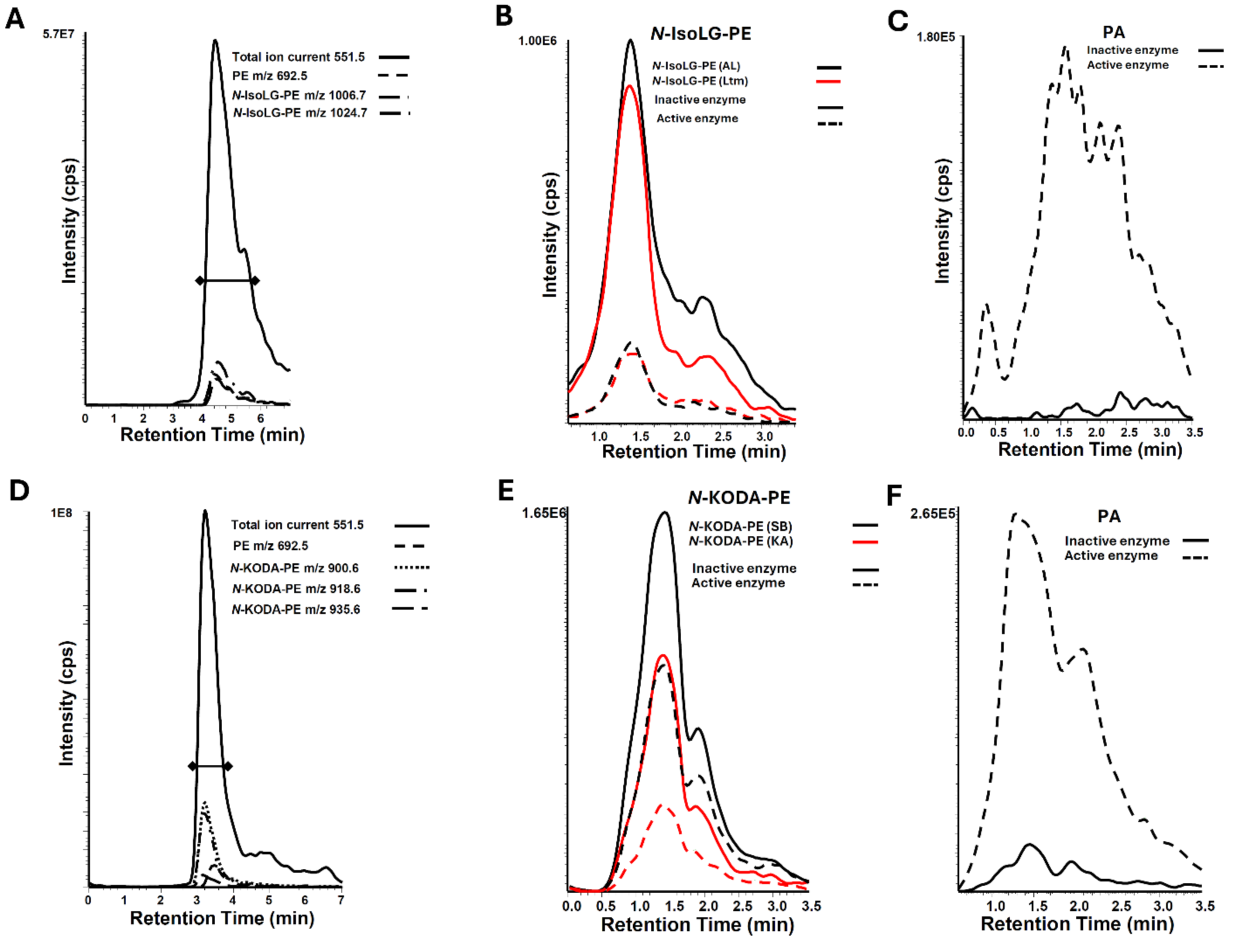


**Supplemental Figure 4. Chromatograms corresponding to LCMS spectra and NAPEPLD hydrolysis of *N*-IsoLG-PE and *N*-KODA-PE in Figure 5.** A) Precursor scanning chromatogram (product ion *m/z 551.5*) of the products of the reaction of IsoLG with PE, resulting in the formation of *N*-IsoLG-PE, including anhydrolactam and lactam species. Solid line represents the total ion current from precursor scan with retention time used for spectra shown in Figure 5A indicated by (♦—♦). Dotted lines represent reconstructed single ion current for major ions found in the spectrum showing *N*-IsoLG-PE spices. B) LC/MS in MRM mode to monitor *N*-IsoLG-PE species and C) phosphatidic acid (PA) as product of the reaction. D) Precursor scanning chromatogram (product ion *m/z 551.5*) of the products of the reaction of KODA with PE, resulting in the formation of *N*-KODA-PE species including Schiff base and ketoamide. Solid line represents the total ion current from precursor scan with retention time used for spectra shown in Figure 5D indicated by (♦—♦). Dotted lines represent reconstructed single ion current for major ions found in the spectrum showing *N*-KODA-PE spices. E) LC/MS in MRM mode to monitor *N*-KODA-PE spices and F) phosphatidic acid (PA) as product of the reaction.

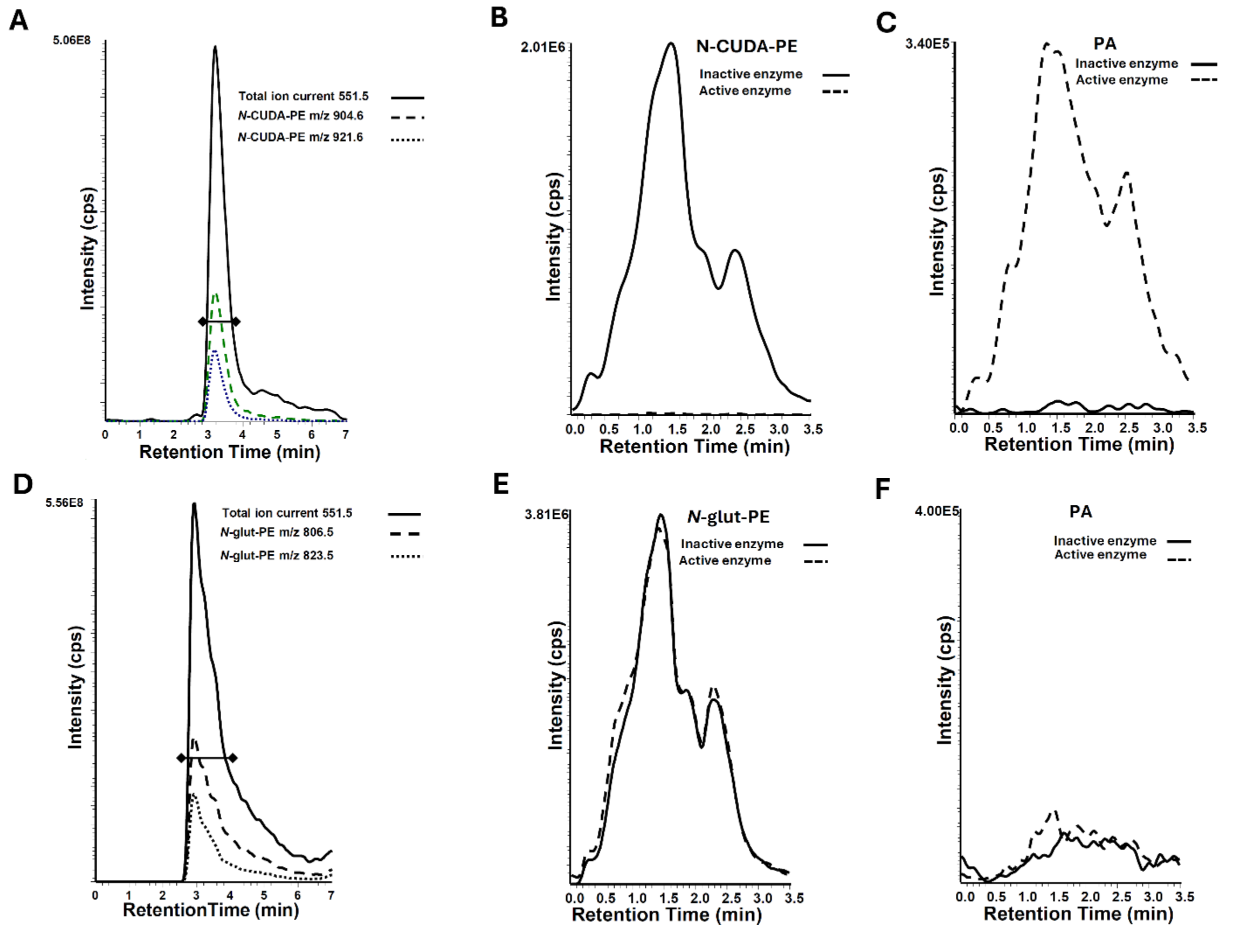


**Supplemental Figure 5. Chromatograms corresponding to LCMS spectra and NAPEPLD hydrolysis of *N*-CUDA-PE and *N*-glutaryl-PE in Figure 6.** A) Precursor scanning chromatogram (product ion *m/z 551.5*) of *N*-CUDA-PE commercial preparation. Solid line represents the total ion current from precursor scan with retention time used for spectra shown in Figure 6A indicated by (♦—♦). Dotted lines represent reconstructed single ion current for major ions found in the spectrum showing *N*-CUDA-PE species. B) LC/MS in MRM mode to monitor *N*-CUDA-PE and C) phosphatidic acid (PA) as product of the reaction. D) Precursor scanning chromatogram of commercially available *N*-glutaryl-PE. Solid line represents the total ion current from precursor scan with retention time used for spectra shown in Figure 6D indicated by (♦—♦). Dotted lines represent reconstructed single ion current for major ions found in the spectrum showing *N*-glutaryl-PE. E) LC/MS in MRM mode to monitor *N*-glutaryl-PE and F) phosphatidic acid (PA) as product of the reaction.
